# Challenges in assessing the roles of nepotism and reciprocity in cooperation networks

**DOI:** 10.1101/372516

**Authors:** Gerald G. Carter, Gabriele Schino, Damien Farine

## Abstract

Nepotism and reciprocity are not mutually exclusive explanations for cooperation, because helping decisions can depend on both kinship cues and past reciprocal help. The importance of these two factors can therefore be difficult to disentangle using observational data. We developed a resampling procedure for inferring the statistical power to detect observational evidence of nepotism and reciprocity. We first applied this procedure to simulated datasets resulting from perfect reciprocity, where the probability and duration of helping events from individual A to B equaled that from B to A. We then assessed how the probability of detecting correlational evidence of reciprocity was influenced by (1) the number of helping observations and (2) varying degrees of simultaneous nepotism. Last, we applied the same analysis to empirical data on food sharing in vampire bats and allogrooming in mandrills and Japanese macaques. We show that at smaller sample sizes, the effect of kinship was easier to detect and the relative role of kinship was overestimated compared to the effect of reciprocal help in both simulated and empirical data, even with data simulating perfect reciprocity and imperfect nepotism. We explain the causes and consequences of this difference in power for detecting the roles of kinship versus reciprocal help. To compare the relative importance of genetic and social relationships, we therefore suggest that researchers measure the relative reliability of both coefficients in the model by plotting these coefficients and their detection probability as a function of sampling effort. We provide R scripts to allow others to do this power analysis with their own datasets.

## Introduction

A propensity for helping others is an adaptive trait when it yields a net return for the actor’s inclusive fitness by increasing direct fitness, indirect fitness, or both (West, Griffin, and Gardner 2007a; Hamilton 1964). Indiscriminate cooperation in a well-mixed population can favor ‘cheating’ whereby less cooperative phenotypes gain the fitness benefits of receiving help from cooperators without paying the same costs of being cooperative, resulting in an ‘evolutionary tragedy of the commons’ (West, Griffin, and Gardner 2007a). Cooperation often takes the form of an investment in specific individuals or types. Individuals can ensure indirect or direct fitness returns on their investments by preferentially helping closer kin, *nepotism* (e.g. Griffin and West 2003; Cornwallis, West, and Griffin 2009) or more cooperative partners, *reciprocity* in the broadest sense (Trivers 1971; e.g. Rutte and Taborsky 2008; Dolivo and Taborsky 2015; Schino and Aureli 2017; Carter 2014; Taborsky, Frommen, and Riehl 2016). Crucially, these strategies can coexist: helping decisions can be influenced by both kinship cues and past experience of reciprocal help. These factors can also interact; for instance, reciprocity can be stronger or weaker among kin than among nonkin (Van Cleve and Akçay 2014; Trivers 1971; Axelrod and Hamilton 1981). Evidence for nepotism and reciprocity can therefore be difficult to disentangle, especially from correlational data (e.g. social networks based on cooperative behaviors such as grooming of food sharing). Indeed, nepotism and reciprocity are causal mechanisms that cannot be directly demonstrated with correlational data. But moreover, we will show how correlational data can lead to incorrect and even opposite conclusions about the relative roles of nepotism and reciprocity caused by asymmetries in the sampling effort needed to accurately estimate kinship versus reciprocal helping. Nepotism can make it difficult to detect correlational evidence for reciprocity even when nepotism is limited and reciprocity is perfect. We discuss the reasons for this and provide a method to help assess the power to detect the effects of kinship and reciprocal help.

For nonhuman animals, a claim of reciprocity is far more contentious than nepotism. Despite growing correlational and experimental evidence, there is still disagreement about the existence and importance of reciprocity outside humans (Clutton-Brock 2009; Carter 2014; Taborsky, Frommen, and Riehl 2016; Schino and Aureli 2017). One reason for this debate is that authors do not agree on what the term ‘reciprocity’ means or should mean (West, Griffin, and Gardner 2007b; Noë 2006; Carter 2014; Bshary and Bergmüller 2008; Lehmann and Keller 2006). Definitions of reciprocity (also called reciprocation, reciprocal altruism, reciprocal cooperation, contingent cooperation, and direct reciprocity) have varied between authors and sub-disciplines (Carter 2014). The original concept of ‘reciprocal altruism’ (Trivers 1971) was quite broad and arguably ambiguous, but subsequent and more narrow definitions of reciprocity restricted its general importance to humans (Carter 2014). For example, the term ‘reciprocity’ has been used to describe: a *broad category* of enforced mutual benefit (analogous to kin selection), a *correlation* between cooperation given and received across dyads or over time (analogous to kin-biased association), a conditional helping *behavior* that causes this correlation (analogous to nepotism), and a specific *psychological mechanism* that might cause this conditional behavior (analogous to phenotype matching) (Carter 2014). For our purposes here, we define *reciprocity* broadly as help given that is influenced by rates of help received (i.e. reciprocal help), where help can involve different behaviors integrated over short or long timespans.

Reciprocity is most evident in controlled experiments where helping rates are immediate responses to past help received, and where partners lack a long-term social relationship (Schweinfurth and Taborsky 2018a; Schweinfurth et al. 2017; Taborsky, Frommen, and Riehl 2016; Dolivo and Taborsky 2015; Rutte and Taborsky 2008). Reciprocity can be harder to test in the context of an enduring social relationship, because this by definition means that the partners will integrate social experiences over longer timespans, and because reciprocal help can take multiple forms, such as allogrooming, food sharing, coalitionary support (Schino and Aureli 2017; Carter 2014; Jaeggi et al. 2013; Seyfarth and Cheney 2012). In the case of such a cooperative relationship, the rates of helping measured by an observer are actually a proxy for a measure of the strength of the underlying causal relationship, rather than the immediate cause of observed reciprocal help. In other words, helping events from A to B should predict helping events from B to A, not because the first event directly caused the second event, but because A and B have a cooperative relationship that causes symmetrical bidirectional helping rates (Schino and Aureli 2010a, 2009). To distinguish between causation and correlation, we use the term *symmetry* for the observed correlation between rates of help given and received—the most common observational evidence for reciprocity. Similarly, we use the term *kinship bias* for the observed correlation between help given and kinship—the most common observational evidence for nepotism.

Semantics aside, reciprocity is also contentious because it is difficult to test, especially in the presence of nepotism. Nepotism is also expected to cause symmetry in helping because kinship is symmetrical; if two sisters often help each other, the help could be pure kin altruism or it could be reciprocal. Whereas kinship bias is considered sufficient evidence for nepotism, symmetry is not widely considered to be sufficient evidence of reciprocity (Carter 2014). A demonstration of reciprocity requires experimentally manipulating helping rates and then measuring a change in reciprocal helping (Schweinfurth and Taborsky 2018a; Dolivo and Taborsky 2015; Rutte and Taborsky 2008; Krams et al. 2013; Krama et al. 2012; Krams et al. 2008; Fruteau et al. 2009). In such tests, kinship can be excluded as a factor by testing only nonkin. However, reciprocity is also expected to play a role in cooperation among kin (Jaeggi and Gurven 2013; Schino and Aureli 2010c; Schweinfurth and Taborsky 2018b; Taborsky et al. 2016; Wilkinson 1984; Wilkinson 1988). An experimental test of both nepotism and reciprocity requires simultaneously manipulating cues to both kinship and past experience of cooperation (Schweinfurth and Taborsky 2018b; Zöttl et al. 2013). The logistical difficulty of such a test explains why the vast majority of evidence for both reciprocity and nepotism is correlational (e.g. Carter and Wilkinson 2013; Schino and Aureli 2010c).

Many studies, especially with primates, have compared the relative effect sizes of reciprocal help and kinship on rates of cooperative behaviors such as grooming or food sharing using correlational data (Jaeggi and Gurven 2013; Schino and Aureli 2010c; Carter and Wilkinson 2013; Koster 2011; Silk et al. 2013; Thomas et al. 2018; Wright et al. 2016; Jaeggi et al. 2016; Engelhardt et al. 2015). A challenge with interpreting this correlational evidence is that the simultaneous effects of nepotism and reciprocity are not equally detectable. Kinship estimates will generally be more precise than estimates of helping rates because of inherent differences in sampling effort. For example, each tissue sample can yield a huge number of genetic markers for assessing dyadic genetic relatedness, but each behavioral sample of watching a group of animals will typically yield only few or no helping events for assessing dyadic helping rates. As a consequence, the more precise estimate of the correlation between kinship and helping can often be over-estimated and detected more easily compared to the less precise estimate of the correlation between help given and received. This means that the presence of nepotism can make simultaneous reciprocity harder to detect.

To assess this idea, we developed a resampling procedure for inferring power to detect both kinship bias and symmetry in mixed-kinship groups. To simulate perfect reciprocity in long-term social bonds, we created data of helping events where individuals based their decisions to help on an unobserved history of past reciprocal help that is perfectly symmetrical within specific pairs. We then systematically changed two variables: (1) the degree of nepotism and (2) the number of observed helping events. Finally, we used permutation and bootstrapping to assess how these two factors interactively influenced the probability of detecting evidence for reciprocity.

To demonstrate the application of our approach to empirical data, we then applied the same permutation and bootstrapping procedures to three datasets where both kinship and reciprocity are suspected to co-exist: allogrooming in female mandrills (*Mandrillus sphinx*), allogrooming in female Japanese macaques (*Macaca fuscata*), and food sharing in female common vampire bats (*Desmodus rotundus*). Mandrills appear to form large groups structured by matriline (Bret et al. 2013; Abernethy, White, and Wickings 2002), and show evidence for reciprocal allogrooming (Schino and Pellegrini 2009) and kin discrimination (Levréro et al. 2015; Charpentier et al. 2007). Japanese macaques are nepotistic, have a despotic social network with a steep dominance hierarchy based largely on maternal kinship, and direct allogrooming to dominant individuals and to consistently preferred partners (Balasubramaniam et al. 2018). Regurgitated food sharing in vampire bats has been a classic example of the possible co-occurrence of reciprocity and nepotism (Wilkinson 1988; Wilkinson 1984).

To test if and how nepotism prevents the detection of evidence for reciprocity, we inferred the power to detect both kinship bias and symmetry in simulated and real datasets of various sizes. To generate slopes and their significance (p-values), we used a permutation test designed to deal with collinearity and non-independence (Dekker, Krackhardt, and Snijders 2007). We plotted the slopes and detection rates for kinship bias and symmetry as a function of sampling effort (sample sizes of observed helping events). These plots show whether the relative roles of kinship and reciprocal help are either remaining ambiguous or becoming clearer with more data. R scripts are available online (Carter et al. 2018) so that others can apply or adapt them to their own kinship and cooperation network data.

## Methods

### Inferring power and precision of symmetry and kinship bias

To infer power, we estimated how estimates of kinship bias (the correlation between kinship and helping given) and symmetry (the correlation between help given and help received) vary with an increasing number of observations (N; note that N is the number of observed helping events, not the number of individuals). To create about 20 equally-spaced values of N, we started at N = 20 and added 5% of the total sample of observed interactions with each next step. For example, a dataset of 500 observations would mean 20 sample size values (N = 20, 45, 70, 95, 120…500 measured events). At each step, we randomly sampled N observations from the total dataset. We sampled with replacement (bootstrapping) to avoid confounding smaller variances at larger samples sizes with smaller variances in our samples. We bootstrapped the datasets 1000 times at each sample size. For instance, at the first step we randomly sampled 20 observations with replacement 1000 times. To analyze the simulated data (described below), we created a different dataset of size N observations by sampling from the given probability distributions 1000 times, rather than bootstrapping a single dataset 1000 times as we did with the empirical data.

For each observed dataset, we extracted the observed coefficients of kinship and reciprocal help from a matrix permutation test: multiple regression quadratic assignment procedure with double semi-partialling (MRQAP-DSP, (Dekker, Krackhardt, and Snijders 2007)). We defined the response variable ‘help’ for individual A to B as the total of duration of help from A to B, divided by the total duration of help received by B for all times where A could have helped B. This measure controls for differences in sampling time, and current situational factors such as need (Farine 2015). ‘Reciprocal help’ for A to B is defined as help from B to A. We applied a log transformation to the empirical allogrooming and food sharing durations because they were lognormal. We z-transformed all variables to obtain standardized beta coefficients, so that an observed coefficient of X for kinship indicates that a one standard deviation increase in kinship predicts an increase of X standard deviations in help.

To calculate p-values for the observed coefficients, we used network-level permutations (Farine 2017) randomizing each input variable independently using the standard approach from the MRQAP-DSP function in the R package ‘asnipe’ (Farine 2013). We used this procedure to generate one null coefficient from a randomized network for each observed coefficient, resulting in 1000 observed and 1000 paired null coefficient values for the two predictors, kinship and for reciprocal help, at each sample size step. At each sample size, we then calculated (1) the mean and 95% confidence interval (CI) for the observed coefficients, which are the observed symmetry and kinship bias estimates, (2) the mean and 95% CI for the null coefficients, which are the symmetry and kinship bias estimates expected under the null hypothesis, (3) the proportion of samples where the observed coefficient was greater than the paired null coefficient, which indicates if the effect is real using all the samples, and (4) the proportion of observed coefficients that were greater than 95% of the expected null coefficients, which indicates the power to detect an effect with one sample of a given size.

### Simulating data with perfect symmetry and 0-100% nepotism

We simulated 500 observations of help among 20 individuals. To simulate correlational outcomes expected from perfect reciprocity, we generated a weighted directed network of symmetrical social bonds, such that the helping rate from A to B was always equal to helping rate of B to A. This symmetrical probability of helping, or social bond strength, could be imagined as representing a history of past unobserved helping interactions in which both individuals helped each other in both directions many times, in which case individuals that did not reciprocate therefore no longer have a strong social bond. To create an event, we then randomly sampled one individual as the actor and selected a remaining individual as the recipient with a probability that was proportional to the social bond strength. The duration of help was also equal to the social bond strength. All observed helping was therefore determined by a symmetrical social bond.

To simulate nepotism as an additional factor, the social bond strength must also correlate with kinship to varying degrees. Nepotism determines the degree to which kinship predicts past reciprocal helping, so we calculated social bond strength (*b*), as a combination of a random kinship value (*r*) and a random non-kinship value (*c*), weighted by a ‘nepotism coefficient’ (*n*), which ranges from 0 to 1:

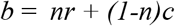

The nepotism coefficient therefore represents the degree to which the social bond strength (and hence the probability of helping) correlates with kinship. For simplicity, we sampled *r* and *c* from uniform distributions, but we obtained similar results from other distributions. Increasing nepotism will increase the observed kinship bias, and we created populations where nepotism equaled either 0, 0.25, 0.5, 0.75, or 1 (Table S1). Finally, we added a step to ensure that all individuals were observed helping at least one other individual.

In sum, these simulations generated an observed set of helping events where individuals could have based their actual helping decisions entirely on the experience of past reciprocal help. However, this reciprocity co-existed across a spectrum of nepotism from 0% nepotism, where helping rates were symmetrical and kinship played no role, to 100% nepotism, where helping rates were symmetrical but social bonds only ever formed among kin, such that the relative causal roles of reciprocal help and kinship are therefore unclear without experimental evidence. There are of course many possible causes of symmetrical helping besides reciprocity. The point of this simulation is to ask: If perfect reciprocity did exist among individuals that were also somewhat nepotistic, how likely are we to detect the evidence for reciprocity or to overestimate the evidence for kinship bias?

### Real datasets

We applied this resampling procedure as a power analysis for three real datasets. Each dataset including a list of cooperative interactions (either grooming or regurgitated food sharing), the duration of the trial (sampling period) during which each occurred, the individuals present during the trial (possible actors and receivers), the actor, the receiver, and the interaction duration. The first two studies were conducted on mandrills and Japanese macaques housed at the Rome Zoo (Bioparco) in Italy. In both studies, kinship was based on maternal pedigrees, all subjects were available as potential grooming partners during the study, and an observer recorded the duration of all female-female grooming episodes involving a focal subject as actor or recipient. The first dataset contained 1703 observations of mandrill allogrooming collected between July 2014 and June 2015 from 10 sexually mature female mandrills in a group that also included two mature males. A past study of six female mandrills from the same captive population found that allogrooming A to B predicted allogrooming B to A, when controlling for kinship (Schino and Aureli 2010c), or when controlling for kinship and rank and excluding recent reciprocal grooming (Schino and Pellegrini 2009).

The second dataset contained 737 observations of macaque allogrooming collected between April and November 1996 from 22 sexually mature female Japanese macaques in a group of 71 that also included mature males and juveniles. Similar to the mandrills, analyses of allogrooming in the same captive population of Japanese macaques found symmetry in female allogrooming, and also found that allogrooming predicted support in social conflicts when controlling for kinship, rank, or time spent in proximity (Schino et al. 2007), and allogrooming was better predicted by kinship than by grooming received (Schino and Aureli 2010c).

The third dataset included 408 regurgitated food-sharing donations among 15 female common vampire bats from previous studies where food sharing was induced by fasting a subject (for details, see Carter and Wilkinson 2015, 2013). Each donation size was estimated by the total seconds that the unfed subject licked the mouth of a fed groupmate. Kinship was estimated using a maternal pedigree and maximum likelihood estimates applied to genotypes of 19 polymorphic microsatellite markers (for details see Carter and Wilkinson 2015). Past analyses of these same data found that food sharing was better predicted by reciprocal sharing than by kinship, when controlling for grooming and donor sex (Carter and Wilkinson 2013a; Carter and Wilkinson 2013b), and this conclusion was supported by later experiments showing that the bats were attracted to the calls of nonkin donors more than nondonor kin (Carter and Wilkinson 2016), and that females that previously fed more nonkin were less affected by the removal of a donor from their food-sharing network (Carter et al. 2017).

### Code availability

Data and R code, including functions to apply the same procedure to other datasets, is available online at the *figshare* data repository (Carter, Schino, and Farine 2018b; Carter, Schino, and Farine 2018a).

## Results

### Simulated data

Nepotism reduced the ability to detect perfect reciprocity in helping. We constructed the simulations such that a perfectly symmetrical social bond determined helping rates at every level of nepotism. At 100% nepotism, symmetry and kinship bias are completely confounded and the relative roles of reciprocal help and kinship cannot be disentangled. At 25% or 50% nepotism, the kinship bias is clearly not as important as the social bond strength. Yet as nepotism increased above zero, the p-values were more likely to incorrectly infer that nepotism was supported by the data while reciprocity was not (Figure 1). The reasons for this can be seen in the plots of the size and precision of the observed and null coefficient estimates with increasing sampling effort (Figure S1-S5). Nepotism increases the correlation between the two predictors: kinship and reciprocal help (Figure S6), but estimates of kinship bias were less variable than the estimates of reciprocal help. At 50% nepotism, kinship biases were often estimated to be larger than reciprocal help, even though the generative probabilities and the actual durations of helping were always perfectly symmetrical (Figure 1). In these scenarios, where we know the real contribution of both kinship and reciprocal help as drivers of helping, we see that kinship bias was consistently overestimated relative to symmetry. Moreover, even with zero nepotism, estimates of symmetry were still underpowered at 500 observations (Figure 1, see also Figure S7).

**Figure 1.**
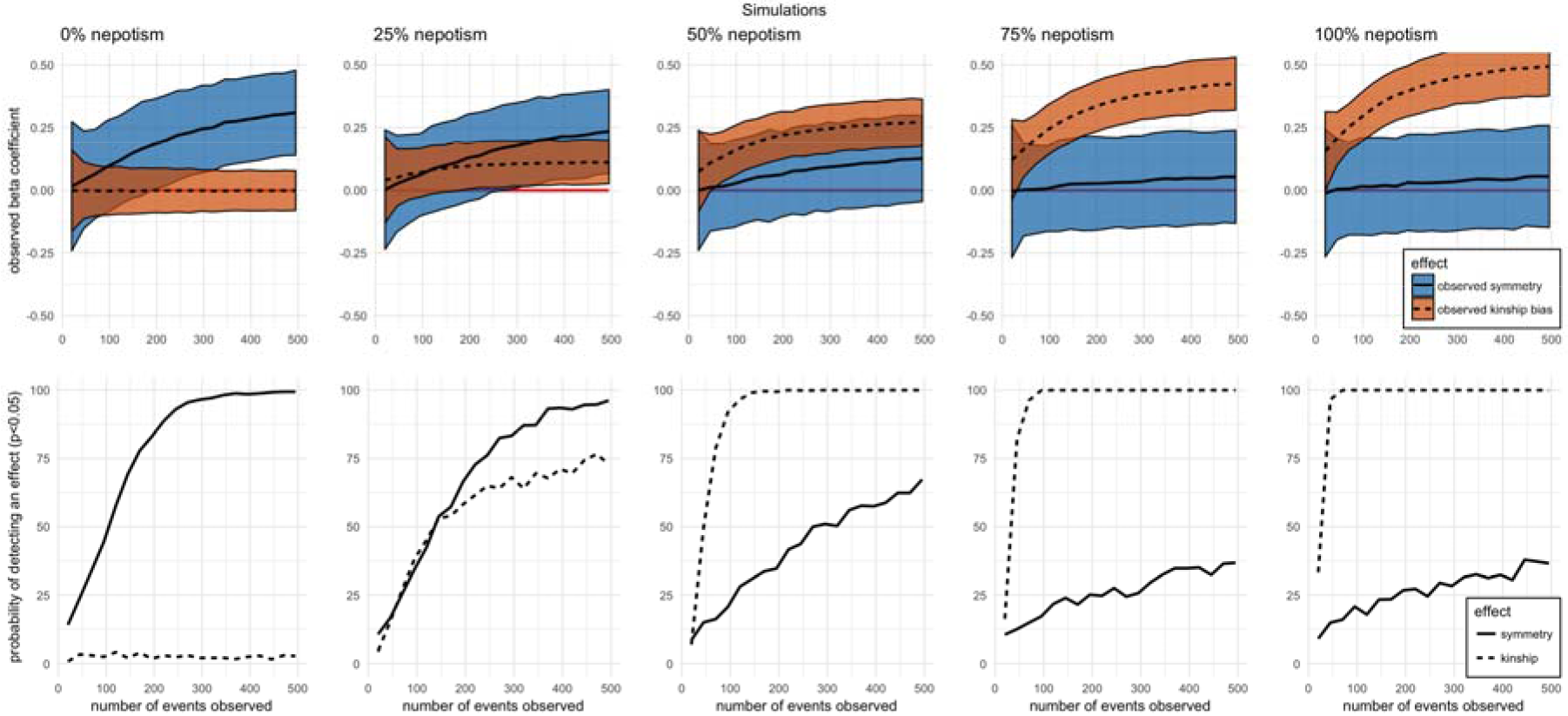
Simulated data: Kinship bias masks perfect symmetry in cooperation networks. The probability and durations of help and reciprocal help are perfectly symmetrical within all dyads, and the degree of simultaneous nepotism increases from left to right. Top panels show the mean and 95% confidence interval of the standardized slope estimates for the effects of helping rate A to B (solid line, purple shading) and kinship between A and B (dotted line, yellow shading) as predictors of helping rate B to A. Bottom panels show the percentage of observed coefficients that were greater than 95% of the coefficients expected based on network permutations. Supplementary Figures S1-S5 in the appendix show plots for the null coefficients and for the probability of the observed coefficients being greater than expected coefficients.

### Real data

Results with empirical data are consistent with expectations from the simulations. For female mandrill allogrooming, symmetry was eventually detected to be significantly greater than kinship bias but this required more than 1250 observations (Figure 2A, 2B). The ability to detect either effect was similar across sampling efforts (Figure 2C, 2D). A sample of 1703 observations provided adequate power to detect both effects, but the relative effect size estimates of symmetry and kinship appear to still be diverging with more observations (Figure 2A), suggesting that the relative contribution of nepotism may still be over-estimated despite this large sample size of helping events.

**Figure 2.**
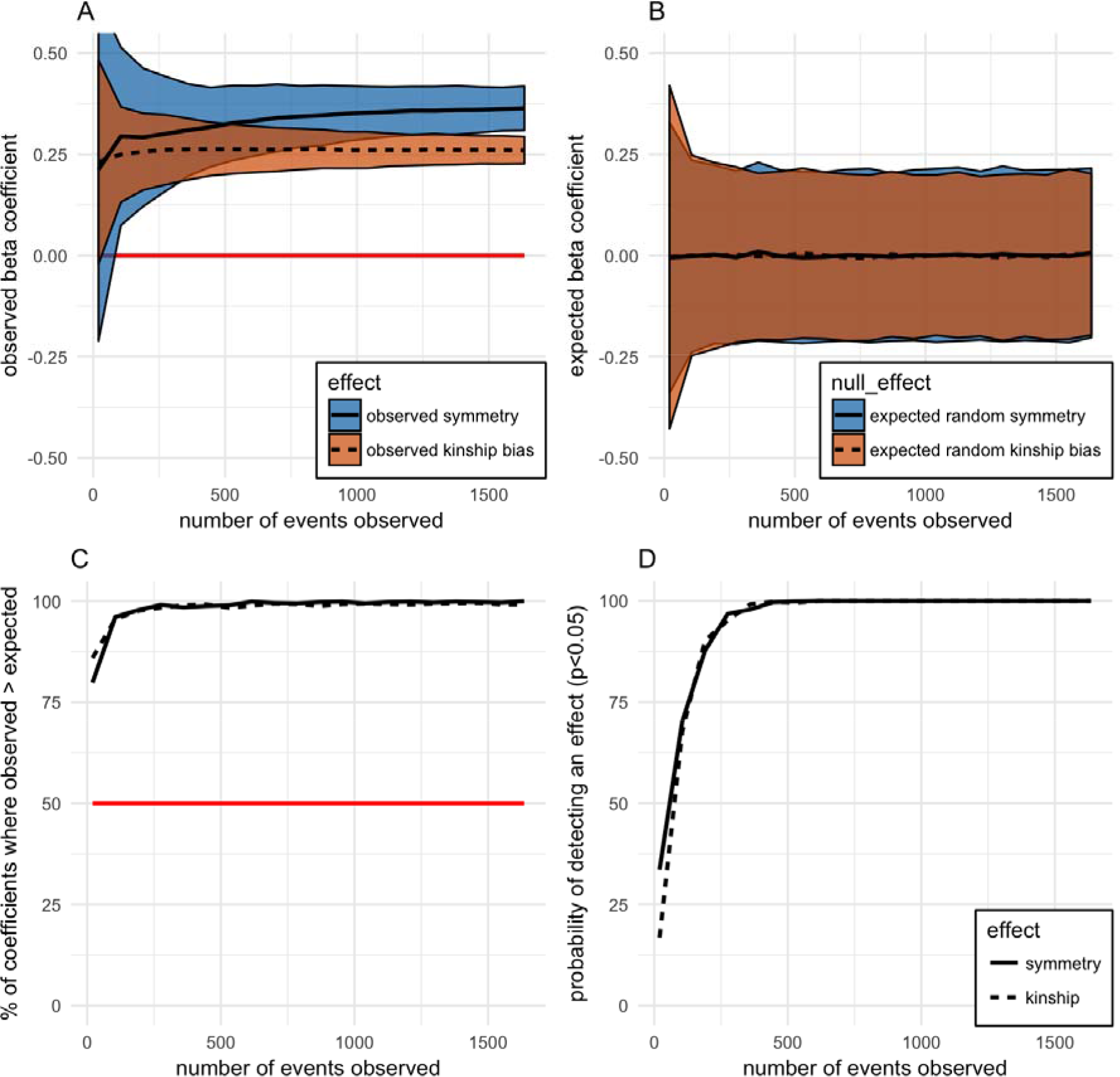
Female mandrills: observed and expected symmetry and kinship bias in allogrooming networks with increased sampling effort. Panel A shows the mean and 95% confidence interval of the standardized slope estimates for the effects of helping rate A to B (solid line, purple shading) and kinship between A and B (dotted line, yellow shading) as predictors of helping rate B to A. Panel B shows the same for the expected null coefficients generated by permutation. Panel C shows the percentage of observed coefficients that are greater than the paired null coefficient generated from the same subsample. If effects are real, then these values should be higher than 50% (red line). Panel D shows the percentage of observed coefficients that are greater than 95% of expected null coefficients for that sample size.

For female macaque allogrooming, where nepotism is quite strong, a sample size of about 60 observations provided enough power to reliably detect a positive kinship bias, but the full set of 737 observations did not provide enough power to reliably detect positive symmetry (i.e. power < 80%, Figure 3). This highlights the combined impacts of greater nepotism and fewer helping events per dyad.

**Figure 3.**
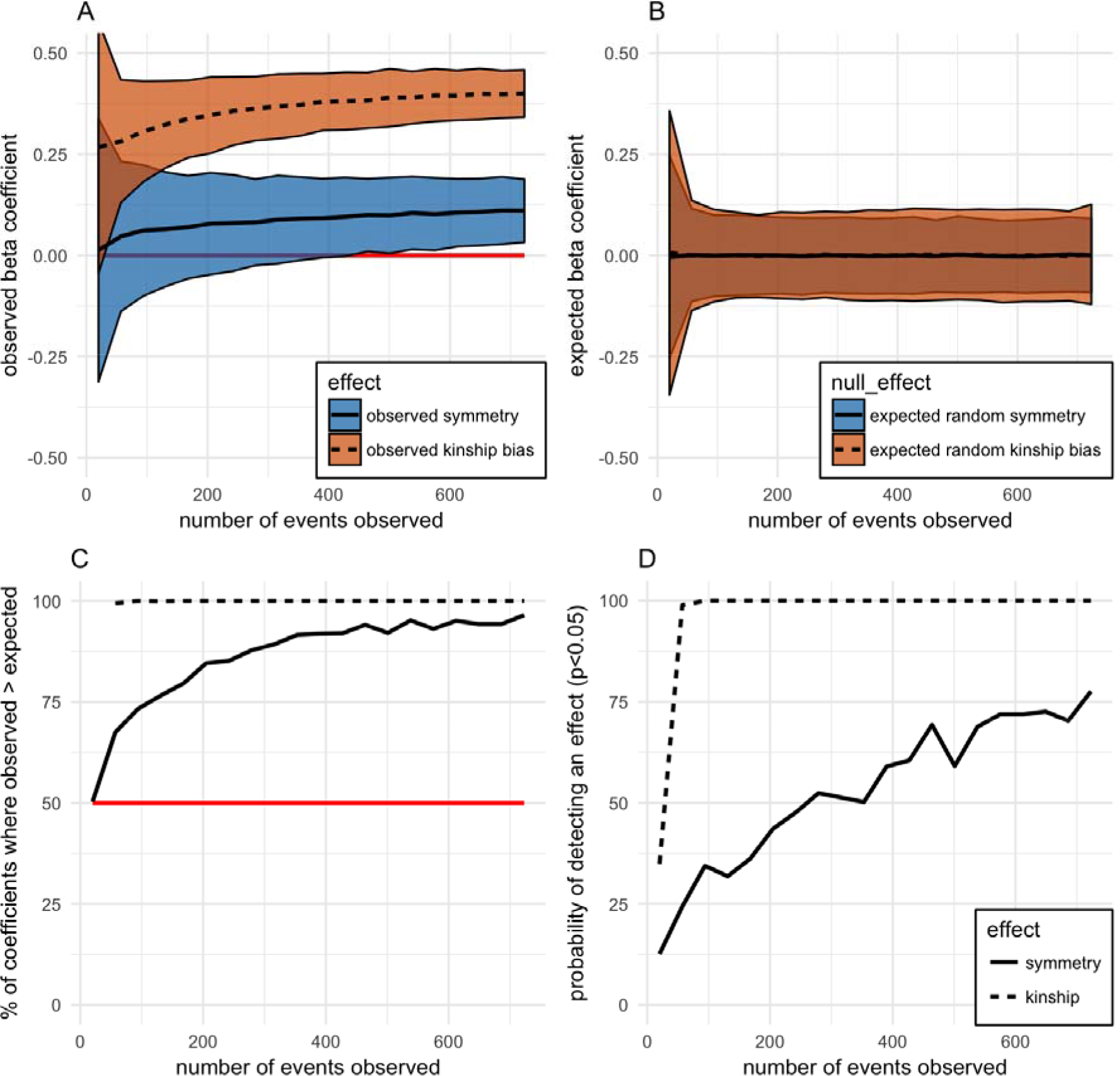
Female Japanese macaques: observed and expected symmetry and kinship bias in food-sharing networks with increased sampling effort. See Figure 2 for explanation of plots.

For food sharing in female vampire bats, kinship bias is detected more reliably than symmetry at all sample sizes (Figure 4). The estimate of kinship bias is relatively stable at about 200 observations, but at 400 observations the estimate of symmetry is still increasing with additional sampling. There is enough power to detect positive kinship bias and symmetry, but power is lacking to reliably estimate the amount of symmetry (Figure 4). Together the plots from our simulations combined with those from the empirical datasets illustrate how and why detecting kinship bias and symmetry requires much less sampling effort than identifying their relative importance.

**Figure 4.**
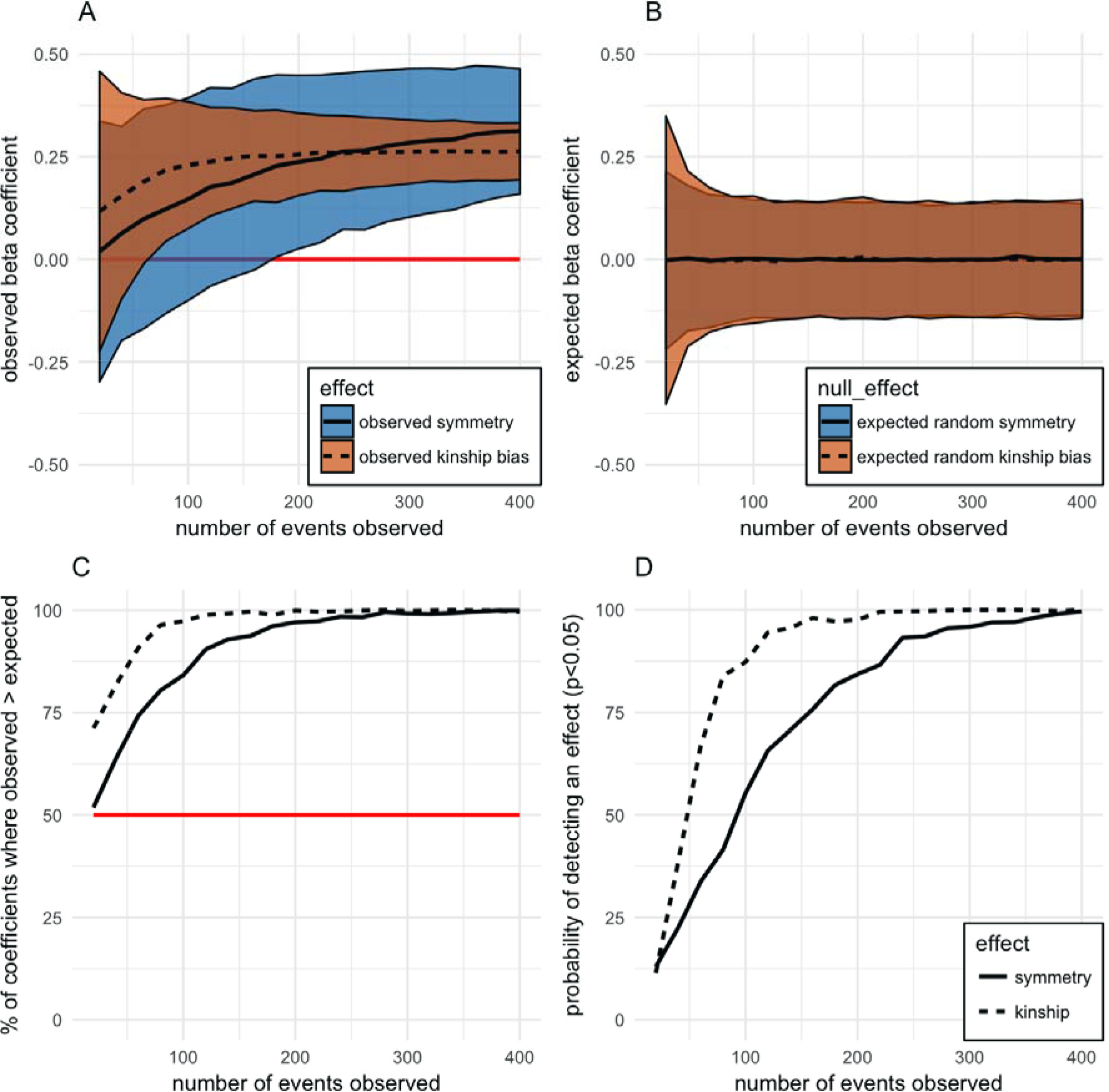
Female vampire bats: observed and expected symmetry and kinship bias in food-sharing networks with increased sampling effort. See Figure 2 for explanation of plots.

## Discussion

### Nepotism can make evidence for reciprocity harder to detect

If kinship and long-term reciprocal help are simultaneous predictors of helping events, then nepotism will often be easier to detect and its role will often be overestimated relative to long-term reciprocal help (or social bond strength). Looking at the simulation results of Figure 1, consider a scenario of 100 helping observations, where individuals are performing perfect reciprocity. At zero nepotism, symmetry (the correlation between help given and received) will be detected (‘statistically significant’) with a 50% probability, and nepotism will have a low rate of false positives. At 25% nepotism, however, a researcher is now equally likely to detect symmetry versus a kinship bias. At 50% nepotism, kinship bias further masks symmetry and the probability of correctly detecting symmetry drops to less than 25% and kinship bias is detected about 90% of the time. At 75% and 100% nepotism, kinship bias is always detected while symmetry is usually not.

Even if both effects are detected, the relative coefficient sizes would give the clear but misleading appearance that helping decisions must be influenced more by kinship than by social experience. In the 100% nepotism scenario, where the symmetry and kinship bias are exactly equal, kinship bias is always detected. By comparison, the same sampling effort in the 0% nepotism scenario gives us only 50% power to detect evidence of reciprocity.

Why does this happen? Estimating a correlation or slope coefficient requires having accurate data on the predictor as well as the response variables. Cases with two highly correlated predictors can lead to imprecise estimates of both coefficients, making each effect harder to detect. More critically, the most precisely estimated predictor, in this case kinship, will often appear to have a relatively larger coefficient and to be more important, regardless of its true causal role. The precision of kinship estimates is independent of the number of helping events observed, whereas symmetry uses the number of helping events to estimate rates of help both given and received.

Genetic relatedness is becoming increasingly easy to measure compared to the history of helping interactions. Pedigrees can be supplemented or replaced by genetic or genomic data, which is becoming cheaper and easier to collect (Städele and Vigilant 2016). The precision of marker-based relatedness estimates is also often estimated by resampling methods. In contrast, the history of cooperation between any two individuals is often unknown and this lack of precision is not always noticed, because there is often confusion about whether each helping event from A to B is just a data point for estimating an underlying cooperative relationship or the direct cause for the next helping event in the other direction (Carter 2014; Schino and Aureli 2017, 2010b; Schino and Aureli 2009; de Waal and Brosnan 2006; Seyfarth and Cheney 2012). In most cases where both reciprocity and nepotism are expected to co-exist (as in the primates and vampire bats examined here), reciprocity is thought to occur in the context of a long-term cooperative relationship. The existence and strength of an underlying cooperative relationship must be estimated by sampling many helping events given and received (e.g. allogrooming, food sharing, or coalitionary support) to calculate dyadic helping rates. When such cooperative interactions are rare or difficult to observe, these dyadic helping rates will be imprecise measures of the true cooperative relationship between the individuals. For example, we might have observed A helping B, but never B helping A, despite the fact that B has helped A in the past, or may do so in the future.

Imprecision in the estimate of helping symmetry will have twice the effect on the ability to find support for reciprocity because it will affect both the predictor (how much B helps A) and the response (how much A helps B), whereas it affects nepotism estimates only once through the response variable (how much A helps B). The sample size of helping observations does not affect the estimates of kinship, but it does determine the accuracy of the dyadic rates of help given and received, so fewer helping observations will directly and greatly impact estimates of symmetry, but not kinship. Even if we observe only one helping event for each dyad, those data could be sufficient to detect and estimate an existing kinship bias, but we could never use the same data to detect evidence for reciprocity. If a cooperative relationship exists, then kinship estimates will generally be more precise than estimates of helping rates (the cooperative relationship).

This asymmetry in precision is compounded in larger study populations. The number of possible dyadic helping rates is almost the square of the number of individuals, n*(n-1). Observing enough dyadic helping interactions to accurately estimate a cooperation network (Farine and Strandburg-Peshkin 2015) is a far greater challenge than estimating pairwise kinship among the same number of individuals. The former requires many unique behavioral samples per individual, whereas the latter might require applying the same panel of genetic markers to one genetic sample per individual. As population size increases, so too does the inequality between the effort required to estimate helping rates versus pairwise kinship across all dyads. As a result, the power to detect helping symmetry relative to kinship bias will almost always decrease with more individuals.

Even when kinship plays a less important role than reciprocal help in determining cooperation, nepotism can essentially overshadow some or all of the evidence of reciprocity, especially with a smaller sample of events. This is important because observations of costly helping events, such as food sharing, are often quite rare. For example, more than 400 hours of focal sampling of wild vampire bats over 26 months of fieldwork yielded 110 observations of regurgitated food sharing and only 21 observations of non-maternal sharing between adults with known association and relatedness (Wilkinson 1984). Researchers studying meat sharing in chimpanzees observed an average of only 1.3 sharing episodes between adult male chimpanzees per successful hunt (Mitani and Watts 2001). When captive capuchin monkeys were separated by mesh barrier and given opportunities to share food, observations of 9896 interactions over food yielded only 18 instances of apparent food sharing and 4 unambiguous cases of one monkey directly giving food to the other (de Waal 1997a). More frequent interactions such as grooming are therefore useful for providing a window into cooperative relationships that may drive more costly helping decisions, but effect sizes based on these most frequent interactions can still be underpowered despite a surprisingly large numbers of helping events (Figure 1).

### How to infer power to detect symmetry and kinship bias

The relative precision of the estimates of kinship bias and symmetry are unclear in a single statistical test. However, as we have shown, one can estimate the relative precision of helping symmetry and kinship bias through resampling methods. Researchers often use resampling to assess the reliability of genetic marker-based estimates of relatedness (Kalinowski et al. 2006; Queller and Goodnight 1989; Wang 2011). Similar methods can be used to create confidence estimates for social relationships (Farine and Strandburg-Peshkin 2015; Sánchez-Tójar et al. 2018).

Our resampling procedure allows one to determine how symmetry and kinship bias respond to increased sampling. For each effect (kinship and reciprocal help), we used subsampling and bootstrapping to generate confidence intervals and trajectories with increasing sampling. By inspecting the resulting plots, one can infer if more power is needed to test a hypothesis about symmetry or kinship bias. For instance, if the coefficients are on a crossing trajectory, then more data are required before drawing any conclusions. If the trajectories are diverging, then more reliable conclusions can be drawn about which predictor is more important, but not by how much. If the trajectories are stable, this suggests that the precision of the estimate would probably not be improved with more sampling.

### Other reasons why reciprocity is hard to detect

We’ve shown that symmetrical helping rates in a nepotistic population can be hard to detect for purely statistical reasons. Several biological factors can also make observational evidence for reciprocity difficult to detect. First, symmetry is more difficult to detect in the timespan of a short-term study because stable relationships are also expected to have a longer timescale for reciprocation (Seyfarth and Cheney 2012; Gomes and Boesch 2011; Gomes and Boesch 2009; Gomes, Mundry, and Boesch 2008; de Waal 1997; Carter and Wilkinson 2013; Jaeggi et al. 2013; Sabbatini et al. 2012; Schino and Aureli 2017). Strongly bonded individuals show dyadic helping rates that are more predicted by a foundation of past events and less predicted by recent events, compared to weakly bonded individuals (de Waal 1997; Seyfarth and Cheney 2012, 1984; Schino and Aureli 2009; Jaeggi et al. 2013). If individuals invest in relationships for long-term benefits, then a strong social bond might only show symmetry over a year but not a month or week. Hence, greater social stability makes it harder to detect helping symmetry over short time periods, whereas lower social stability makes it harder to detect symmetry over long time periods. The relative importance of recent events versus longer-term past experience in driving decisions to cooperate is an open question for most behaviors and species.

Second, relationships are not completely stable and can change during the course of a study. This is generally true for any measure of a social relationship such as dominance or association rate. Social decisions are also influenced by immediate costs and benefits (Newton-Fisher & Kaburu, 2017). For instance, one study reported that an alpha male chimpanzee was attacked and killed by his most strongly affiliated and most frequent grooming partner, that many observers might label as a ‘friend’ (Kaburu, Inoue, & Newton-Fisher, 2013). Social relationships can not only change suddenly, but often asynchronously across dyads.

Third, in addition to the helping rate, the relative importance of reciprocity and nepotism for determining the helping rate can change over time. For example, the care invested by a mother in her two daughters when they are young might be 100% nepotistic and 0% reciprocal, with equal helping allocated to each daughter. However, when her daughters become adults, the mother’s investment might also be influenced by each daughter’s reciprocal investment in her, and she may have a stronger relationship with one daughter over another.

Fourth, reciprocity can be more difficult to detect than nepotism because cooperative returns can take different forms (Fruteau et al. 2009; de Waal and Berger 2000; de Waal 1997; Gomes and Boesch 2011; Gomes and Boesch 2009; Seyfarth and Cheney 1984; Borgeaud and Bshary 2015). If each decision to help is based on a weighted sum of different forms of past help, then any one form of help might not show much symmetry even if the relationship is balanced when all forms of help are considered. The notion of reciprocity by ‘emotional book-keeping’ (Schino and Aureli 2009), implies that ‘grooming on Tuesday can create an emotional bond that causes meat sharing on Saturday afternoon’ (p. 167, Seyfarth and Cheney 2012). Asymmetries in any one service are expected if subordinates groom more dominant individuals in exchange for tolerance (Borgeaud and Bshary 2015; Tiddi et al. 2011; Port, Clough, and Kappeler 2009; Ventura et al. 2006), or if individuals adjust their grooming rates based on the ability of partners to provide food relative to others (Fruteau et al. 2009). Such asymmetries play a key role in biological market theory (Noë and Hammerstein 1995, 1994) but pose a problem for simple ‘tit-for-tat’ models of reciprocity.

Finally, reciprocity and nepotism might interact: the degree of contingency in a reciprocal relationship may be more or less strict among kin. A negative interaction between contingency and kinship was observed in cooperatively breeding cichlids where dominants share their nests with subordinate helpers that must ‘pay-to-stay’, but subordinates nonkin help more than kin because dominants tolerate subordinate kin regardless of their degree of alloparental care (Zöttl et al. 2013). Such differences in reciprocity between kin and nonkin may also exist in primate long-term cooperative relationships, but they can only be detected through repeated manipulations of helping among both kin and nonkin.

### Experimental evidence for simultaneous nepotism and reciprocity

To test the causal roles of reciprocity and nepotism, experiments must manipulate both the helping history and kinship cues that influence decisions to help. To our knowledge, this has only been accomplished once using an experimental paradigm where rats are trained to understand how to pull a bar to deliver a food reward to a partner rat. In a series of experiments, reciprocity was evident because decisions to pull for a partner were influenced by factors such as past food received or allogrooming received from the partner (Rutte and Taborsky 2008; Schweinfurth and Taborsky 2018a). To test for a simultaneous kinship effect, outbred wild-type male rats were separated from littermates, housed with non-kin, tested for an ability to recognize kin, and then tested in the same food-pulling task with partners that varied in both their past reciprocal help and kinship (Schweinfurth and Taborsky 2018b). The rats demonstrated kin discrimination by preferring to associate with unfamiliar kin over unfamiliar nonkin, but they did not show nepotism in the food pulling task; kinship did not increase food pulling nor did it change the symmetry of reciprocal pulling rates (Schweinfurth and Taborsky 2018b).

### When is nepotism harder to detect than symmetry?

There are also several conditions under which nepotism is unlikely to be detected relative to reciprocity, such as when kinship estimates are inaccurate or when there is insufficient variation in kinship among dyads (Csilléry et al. 2006). Although genetic and genomic data is becoming cheaper, easier, and more available (Städele and Vigilant 2016), kinship estimates based on genetic data can still be quite imprecise (Csilléry et al. 2006; Pemberton 2008; van Horn, Altmann, and Alberts 2008). Second, relatedness estimates become severely biased using allele frequencies calculated from only a few animals (Wang 2017). If genetic samples are used to score relatedness in a small subset of individuals, it is crucial to calculate the baseline population allele frequencies from a much larger sample. In studies using pedigrees based on births, ‘kinship’ is actually maternal kinship. These estimates may be ecologically valid if the animals themselves cannot recognize paternal kin, but increasing evidence suggests that some primates for instance can recognize unfamiliar paternal kin (Levréro et al. 2015; Charpentier et al. 2007).

Although helping rates are less precise than kinship estimates, association rates might often be more precise than kinship estimates if dyadic association rates are based on automated methods that can involve many thousands of observations (Aplin et al. 2015; Alarcón - Nieto et al. 2018). In such cases, the social networks could be described more accurately than the genetic relationships, and association will be easier to detect than kinship as a predictor of cooperation.

### Quantifying if and how nepotism contributes to symmetrical helping in network data

If both reciprocity and nepotism exist, decomposing the inclusive fitness benefits of a cooperative trait into the relative fitness and indirect fitness components is implausible using empirical observational data. On the other hand, both controlled experiments and some observational analyses can help to identify the relative roles of different proximate mechanisms. Experiments can directly identify the relative importance of different conflicting cues used to make helping decisions (Schweinfurth and Taborsky 2018b). Observational studies can also play an important role in looking at what proximate factors best predict cooperation in nature by investigating how cooperation network structures arise. For example, consider two hypotheses. In the first, kinship determines proximity, which then determines symmetrical grooming. In the second, individuals associate in space independent of kinship, but they preferentially groom their kin. Recent developments in social network analysis (Farine and Whitehead 2015) and null models (Farine 2017) can provide potentially useful tools for distinguishing between such scenarios by constructing different mechanistic networks (VanderWaal et al. 2014; Ilany and Akçay 2016) or by constructing different null models that allow different aspects of associations to vary (Farine et al. 2015). Linking these models to tests of symmetry and kinship bias could yield greater insights into whether kinship or past experience shape patterns of helping directly or via more simple processes (Puga-Gonzalez, Hoscheid, and Hemelrijk 2015).

Ultimately, patterns generated by reciprocal helping should have a temporal signature in that helping given should reflect some degree of helping received in the past. However, as we noted above, a major challenge is determining over what timeframe reciprocity takes place in a stable cooperative relationship. Few studies to date are likely to have a sufficiently complete dataset of helping behaviors to use temporal analyses. However, once such data are available, temporal social network analysis (Pinter-Wollman et al. 2013; Blonder et al. 2012; Farine 2018) could provide useful tools for investigating these topics.

### Practical recommendations

When testing simultaneously for evidence of nepotism and reciprocity in cooperation networks, researchers should be aware of the necessity to estimate the reliability of estimates of both kinship and helping rates (the network edge weights) before comparing their relative importance. The most obvious way to improve inferences is to collect more interactions and to estimate kinship using more pedigree or genetic data. Adding more individuals, however, cannot compensate for a lack of repeated measures of the same individuals, which is what determines the precision of network edge weights (Farine & Strandburg-Peshkin 2015). One way to increase such repeated measures is to induce acts of cooperation. For example allogrooming can be induced by applying substances to the fur (Hemelrijk 1994; Schweinfurth, Stieger, and Taborsky 2017), cooperative mobbing can be induced with fake predators (Krams et al. 2013; Krama et al. 2012; Krams et al. 2010; Krams et al. 2008), food sharing can be induced by fasting individuals (Carter and Wilkinson 2013; Wilkinson 1984) or by creating opportunities to provide (Silk et al. 2013).

We used network permutations which account for the network structure and hold the total help given and received by each individual constant. However, network permutations do not account for biased sampling, so the helping rates (network edge weights) must take into account the relative opportunity for individuals to help each other. We accomplished this by defining edge weights as the proportion of help received from individual X divided by the total help received from all other individuals that could have otherwise come from individual X because X was present at the time. Another possibility is to define edges as the help from X over the opportunity for X to help. If helping events are scored as yes/no events, then an even more rigorous approach is to use pre-network permutations (Farine 2017), where the helping acts in the dataset are permuted across individuals present at the time, rather than permuting the helping rates in the network. Pre-network permutations allow for precise control over the null hypothesis by swapping within time periods or locations, and also control for biased sampling; however, they are most appropriate when the helping events are binary (0/1) and hence interchangeable. In conclusion, due to differences in the ease of detecting symmetry and kinship bias, it is useful to assess the reliability of each effect as a function of sampling effort. We provide R code (Carter et al. 2018) to produce plots that allow one to assess the relative power for detecting evidence of nepotism and reciprocity in simulated datasets or in a given dataset of helping observations in humans or nonhuman animals.

## Acknowledgements

We thank T Chen, I Hamilton, J Massen, and an anonymous reviewer for constructive feedback which improved the manuscript. We thank the Rome Zoo (Bioparco) for allowing us to study their Japanese macaque and mandrill colonies, and Francesca Lasio and Raffaella Ventura for help with the data collection. GGC was supported by fellowships from the Smithsonian Institute and the Humboldt Foundation. DRF was funded by the Max Planck Society.

## Appendix

Gerald G. Carter, Gabriele Schino, Damien Farine

**R code can be found here:** https://doi.org/10.6084/m9.figshare.6072272.v2

**Datasets can be found here:** https://doi.org/10.6084/m9.figshare.6072254.v2

The R function “infer_power_reciprocity_kinship” uses the following arguments:

- **individuals:** a vector with unique IDs for all individuals in the study
- **observations:** a data frame with columns: actor, receiver, duration, possible.actors.list
- **possible.actors:** a matrix where each column contains a list of possible actors for each observation. The number of rows is the max number of possible actors.
- **relatedness:** a matrix of relatedness between all individuals - row and column orders must match the order of individuals
- **nreps:** number of repeated subsamples at each sample size
- **jumps:** number of observations to increment each sample size step (default = 20). First sample size is always 20.
- **simulate_data**: if set to “TRUE”, the function will simulate data. If set to “FALSE” then the function will use actual data provided by arguments above.
- **nepotism** (only used with simulated data): degree of nepotism (0 to 1)
- **N** (only used with simulated data): number of simulated individuals
- **n.events** (only used with simulated data): number of simulated helping events

Note: Depending on the number of observations and replications, this function can take several hours to run.

---

**Table S1.** Mean correlation between social bond strength and kinship in simulated data. Values based on 10,000 simulations. Social bond strength is always symmetric and determines probability and duration of observed helping events.

| Nepotism | Mean Pearson’s correlation (0.025 – 0.975 quantiles) |
| --- | --- |
| 0% | 0 (-0.14—0.14) |
| 25% | 0.32 (0.19—0.44) |
| 50% | 0.71 (0.64—0.76) |
| 75% | 0.95 (0.94—0.96) |
| 100% | 1 (1—1) |

**Figure S1.**
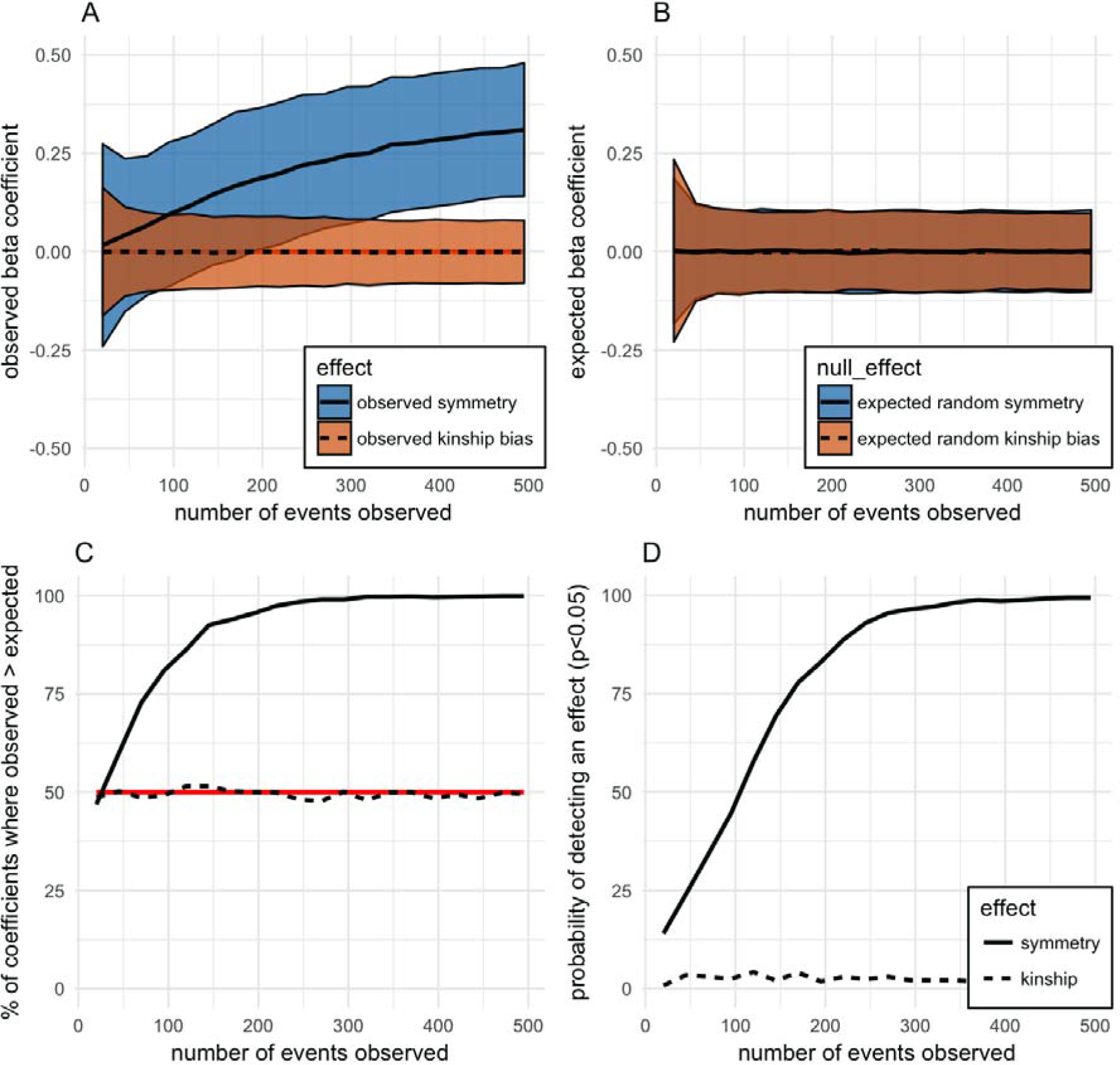
Power analysis for simulation of symmetrical helping with 0% nepotism. Panel A shows the mean and 95% confidence interval of the standardized slope estimates for the effects of helping rate A to B (solid line, purple shading) and kinship between A and B (dotted line, yellow shading) as predictors of helping rate B to A. Panel B shows the same for the expected null coefficients generated by permutation. Panel C shows the percentage of observed coefficients that are greater than the paired null coefficient generated from the same subsample. If effects are real, then these values should be higher than 50% (red line). Panel D shows the percentage of observed coefficients that are greater than 95% of expected null coefficients for that sample size.

**Figure S2.**
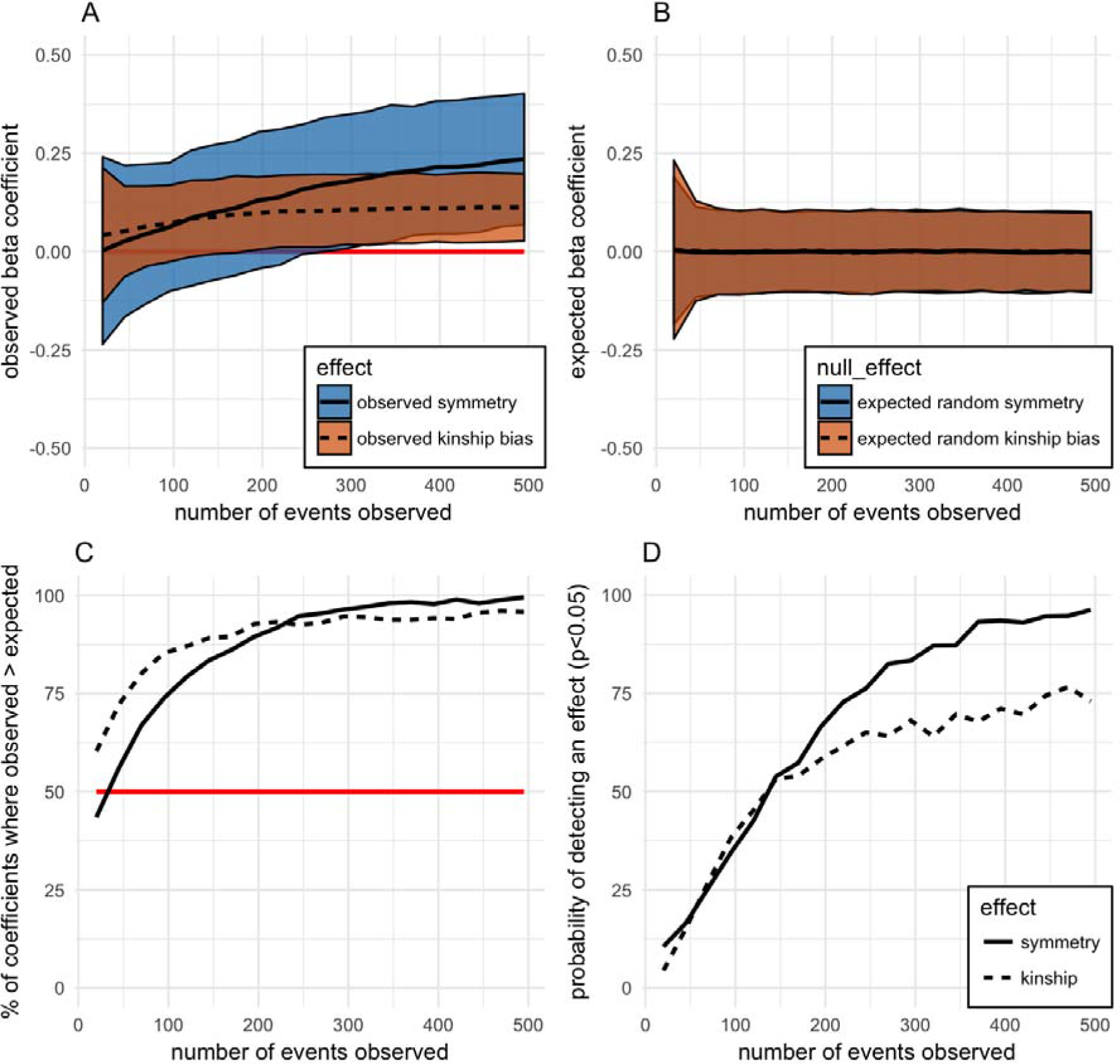
Power analysis for simulation of symmetrical helping with 25% nepotism. See plot Figure S2 for explanation.

**Figure S3.**
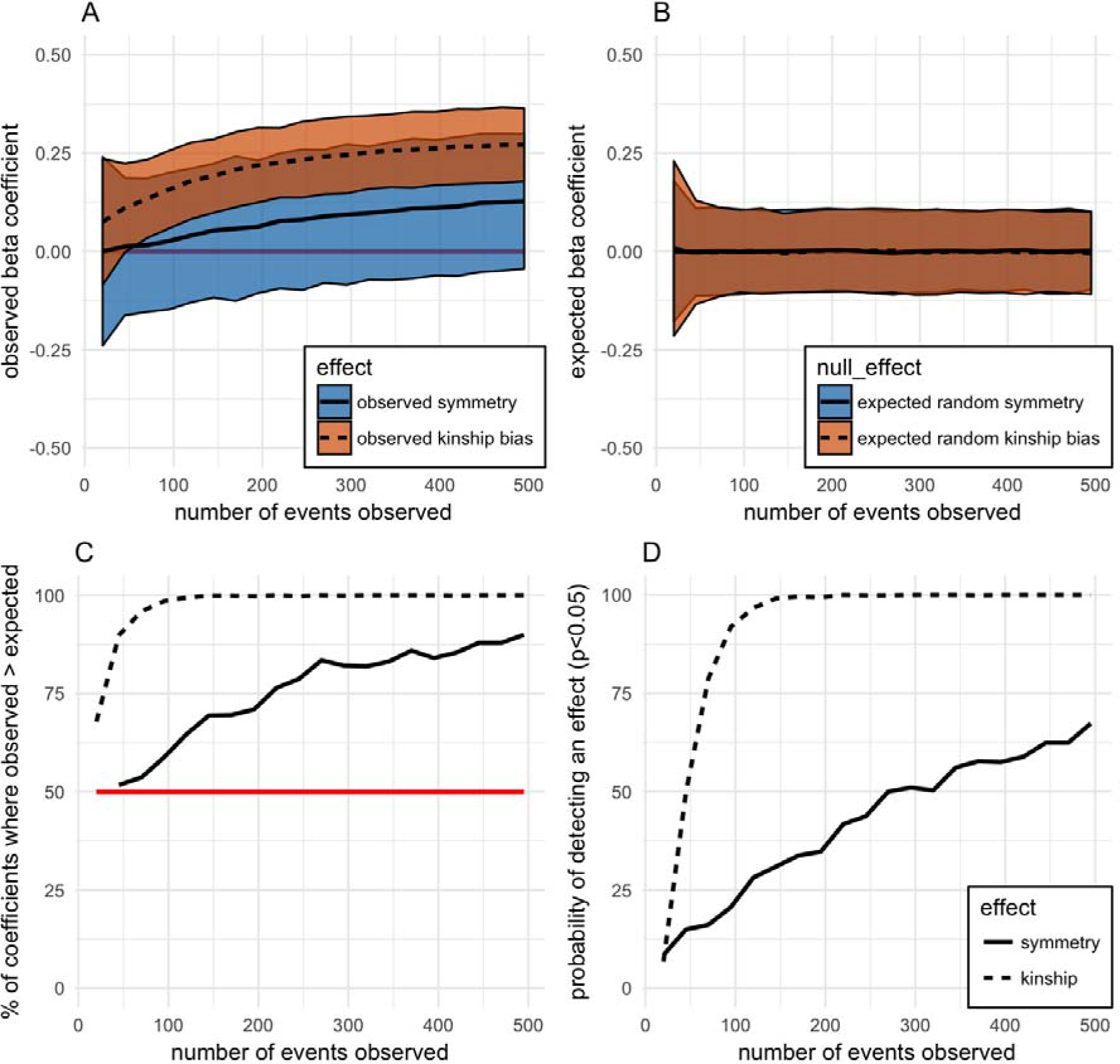
Power analysis for simulation of symmetrical helping with 50% nepotism. See plot Figure S3 for explanation.

**Figure S4.**
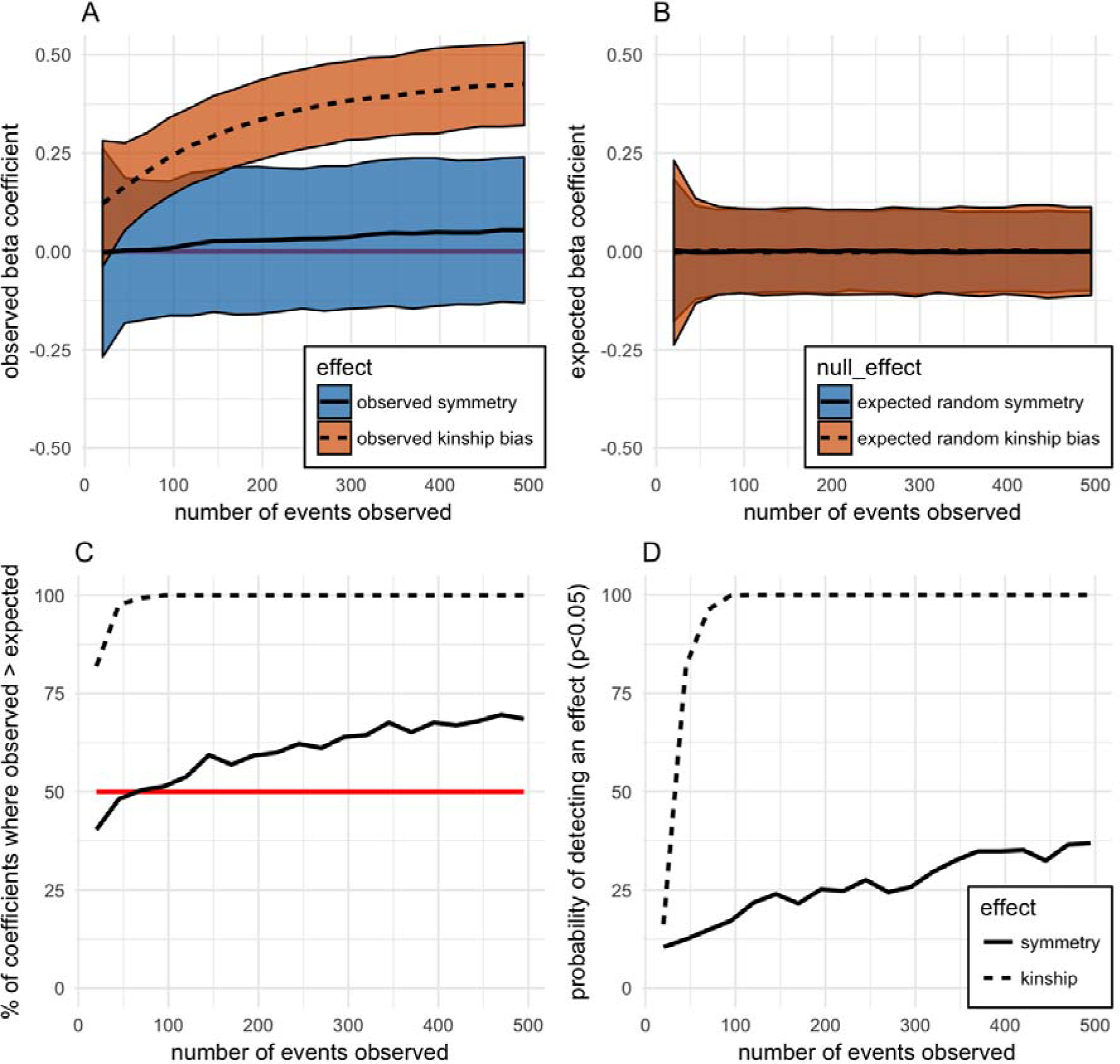
Power analysis for simulation of symmetrical helping with 75% nepotism. See plot Figure S4 for explanation.

**Figure S5.**
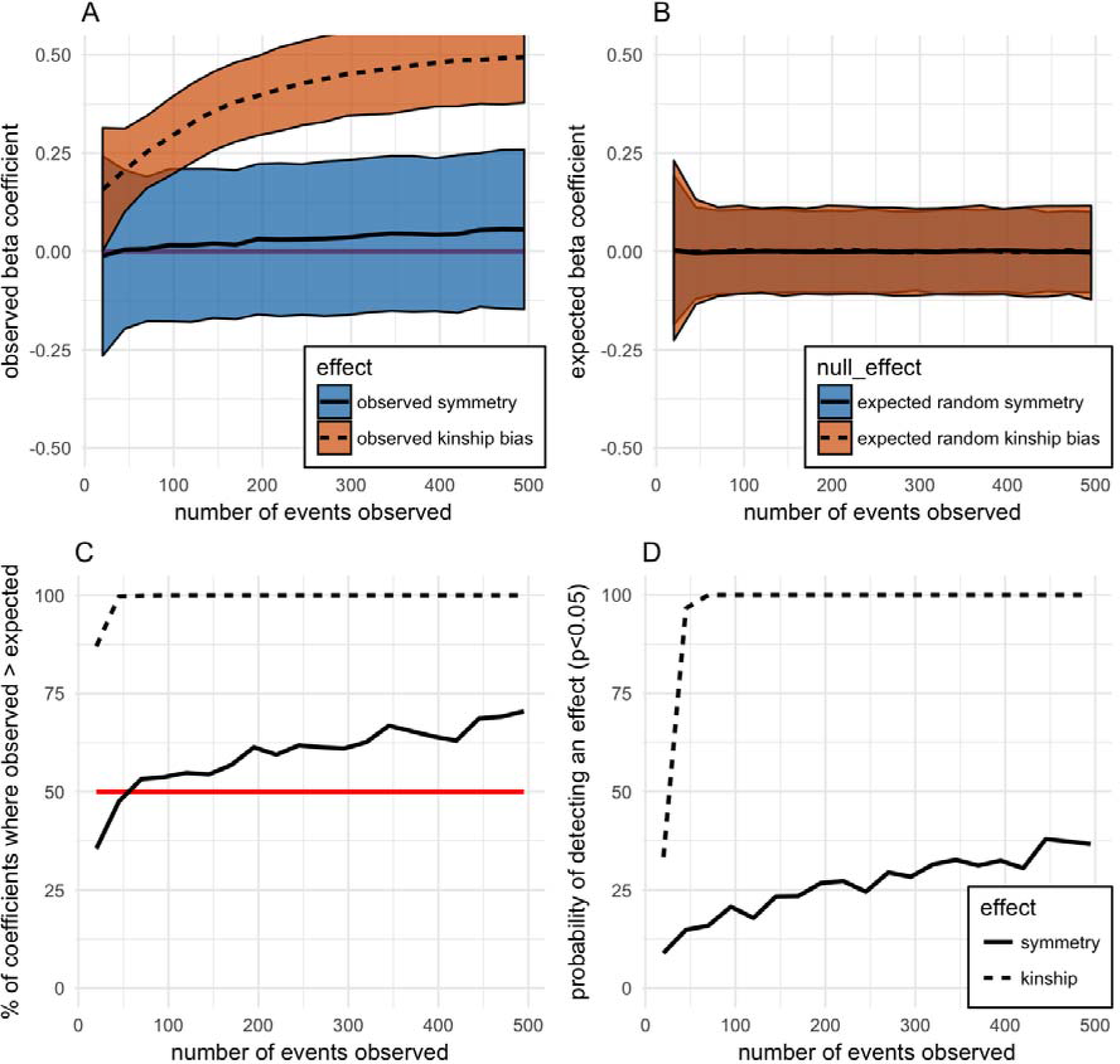
Power analysis for simulation of symmetrical helping with 100% nepotism. See plot Figure S5 for explanation.

**Figure S6.**
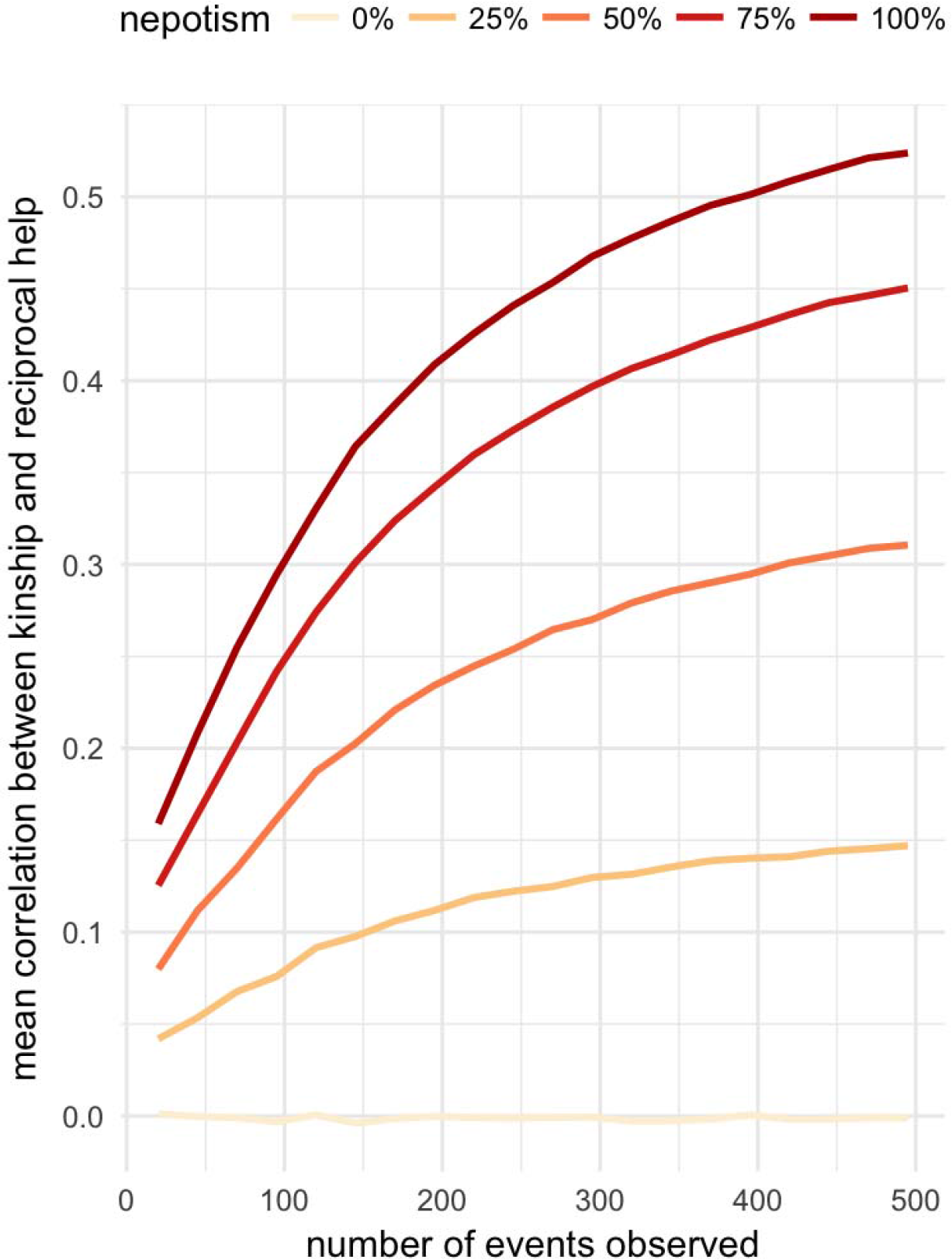
Collinearity. Mean Pearson’s R correlation between the predictors kinship and reciprocal help for each degree of nepotism with increased sampling effort.

**Figure S7.**
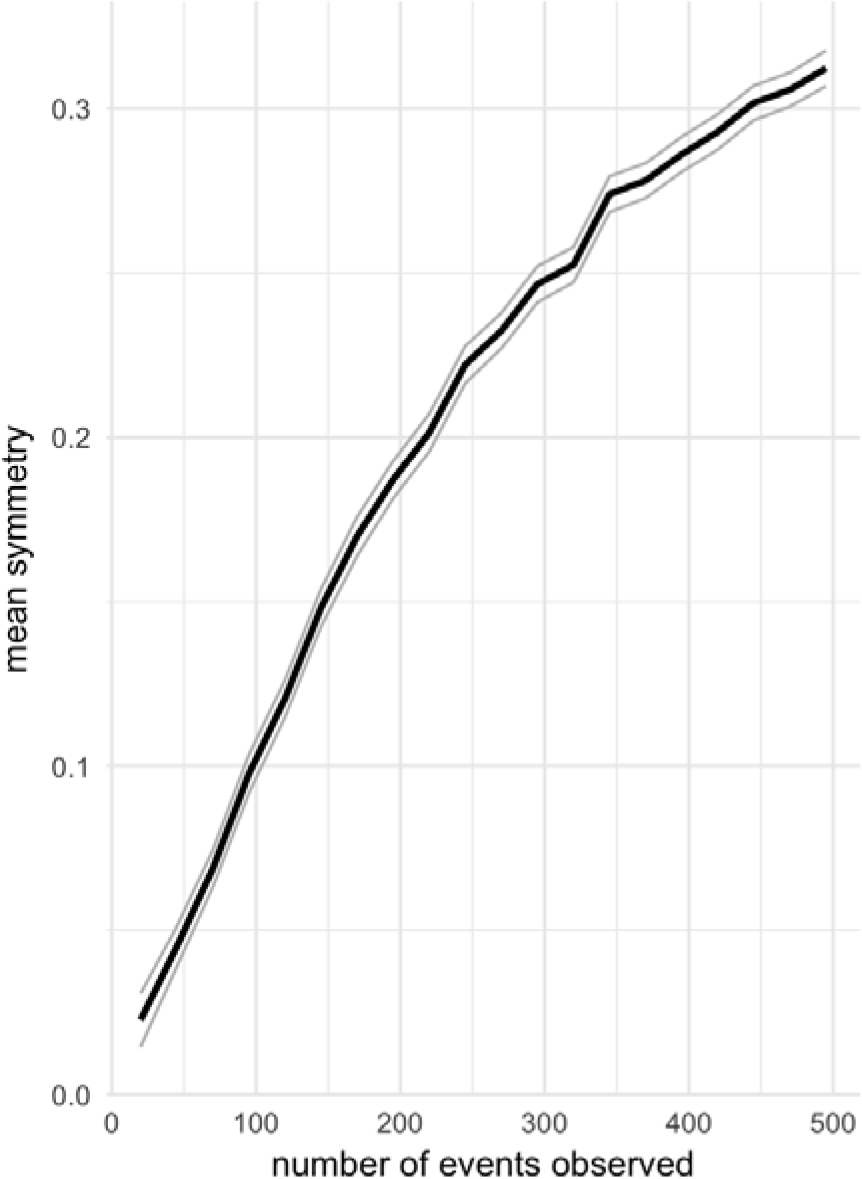
Symmetry with increased sampling effort expected with perfect reciprocity and zero nepotism. Plot shows mean (black) and 95% confidence interval (grey) of Pearson’s correlation between help given and received.

